# Computational and experimental hunt for expansion prone tandem CNG repeats in human genomes

**DOI:** 10.1101/2023.05.03.539188

**Authors:** Varun Suroliya, Bharathram Uppili, Manish Kumar, Vineet Jha, Achal K Srivastava, Mohammed Faruq

## Abstract

**Introduction:** Spinocerebellar ataxias (SCA) are a group of clinically and genetically heterogeneous neurodegenerative disorders. Tandem repeat expansion is the pathogenic mutation in most of SCA cases. The pathophysiology of SCAs is still not completely defined due to the lack of genetic mutation in around 50% of cases worldwide. These uncharacterized cases must be genetically diagnosed for a better understanding and future treatment purposes. In this study, we tried a combination of computational and experimental approaches to find out novel CNG repeat loci that may be associated with SCAs.

**Methodology:** We investigate the human reference genome (hg-37) to find out all the possible CNG repeats present in more than 3 continuous uninterrupted units and annotated their functional locations. For experiment purposes, we targeted 52 loci from exonic and UTR regions and screened them in our 100 control samples through fragment analysis to know their polymorphic status. All the highly polymorphic loci were further investigated in 100 patient samples to know any large repeat expansion.

**Results:** There are 15069 CNG repeat loci present in the human genome. After the examination of 52 loci in the control samples, 19 loci showed a highly polymorphic repeat pattern and were screened in patients. The 1000 genome different population data also suggested the polymorphic status in the available 15 loci data. From the GTEx database, 18 loci proposed the expression in the brain, suggesting any variation in these genes may cause neurological disorders.

**Conclusion:** We tried a different kind of approach to find out tandem repeat expansion mutation in a cost-effective manner. Although we were unable to identify any disease-causing mutation in our patient cohort recently, various studies from different populations of the world have vouched for the genetic changes in these genes like GLS, RAI1, GIPC1, and CNKSR2 resulted in neurological disorders. Moreover, publications on the same identified repeats in GIPC1 for OPMD and GLS for ataxia with developmental delay, confirm the hypothesis to evaluate these repeat loci in different populations and other neurological disorders to identify novel targets.

## Introduction

Spinocerebellar ataxia is a group of neurodegenerative disorders. The most common genetic mutation in SCAs is repeat expansion in the coding or noncoding gene region. To date, more than 40 genetic repeat loci were reported associated with ataxic dysfunctions still 20-60% of cases are genetically unclassified worldwide (1) (Fig-1).

**Fig 1.**
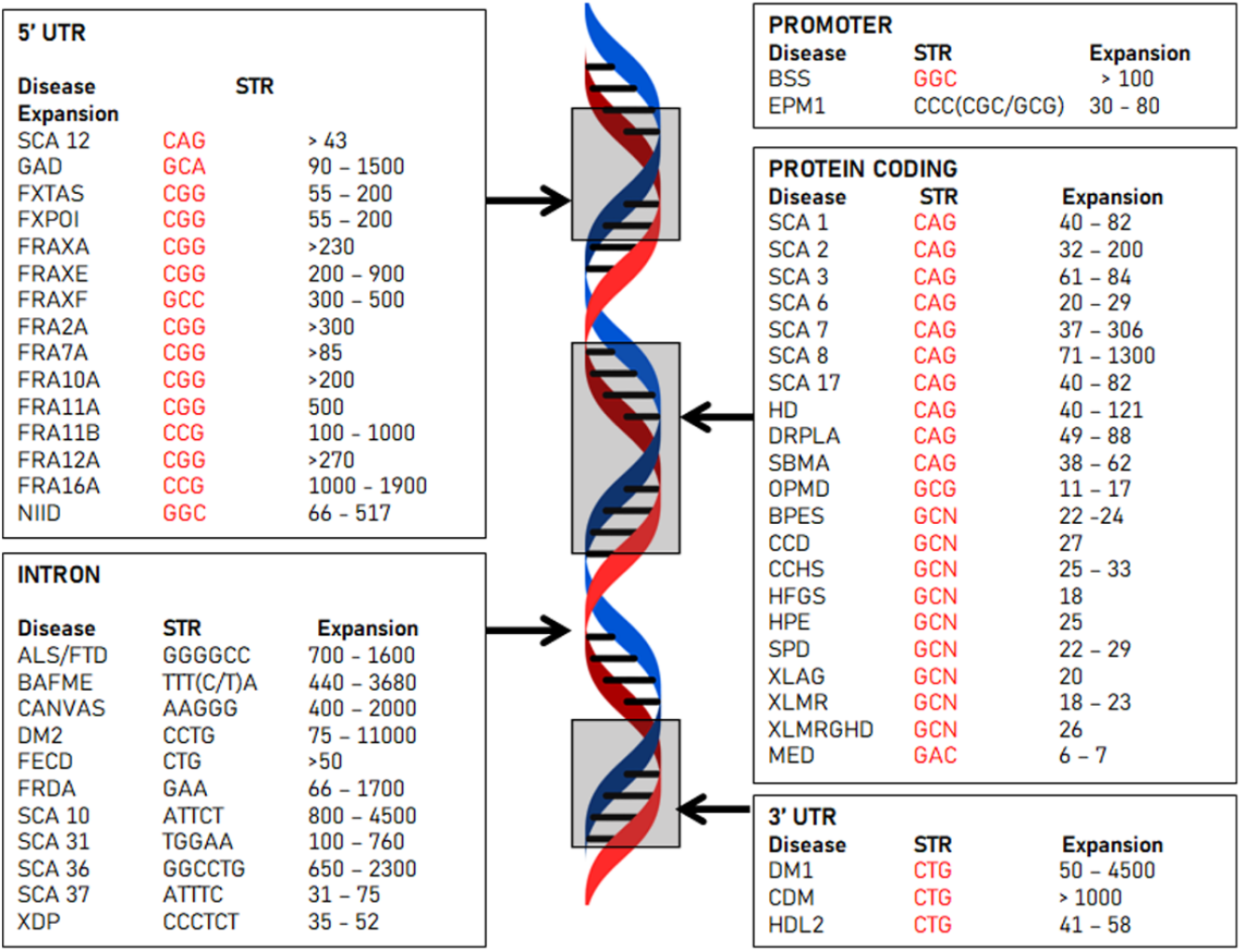
Illustration of short tandem repeat [STR] expansion disorders on the basis of their functional region of the genome together with the disorders and pathogenic expansion range.

The available literature, various databases as well as our in-house data explain the role of CNG as the most prevalent cause of SCA in autosomal dominant inheriting patients. Thus, prioritization of such loci becomes essential for successful candidate gene identification. The majority of the SCA subtypes are geographical region specific and different countries are dealing with specific subtypes. In the north Indian SCA cohort, SCA 2 and SCA 12 are the most common autosomal dominant cerebellar ataxias (ADCAs) and FRDA in the autosomal recessive cerebellar ataxias (ARCAs) sub-type (2), whereas SCA 3 is the most common subtype worldwide (3). In the Indian spinocerebellar ataxia patients, ∼ 60% are still genetically uncharacterized. This raises the possibility of other trinucleotide repeat expansion might be responsible for these cases.

Identification of novel repeat loci is a very difficult task due to the rarity of the disease, clinical heterogeneity in symptoms, difficulty to amplify, or even finding from short-read sequencing data, high cost of long-read sequencing, and many more.

In 2004 Pandey et al. tried a different type of approach and computationally reviewed the entire CAG repeats in the genome and identified two CAG loci as putative candidates for SCA disorder (4). In this study, we used a combination of computational as well as genetic approaches to identify possible disease-causing unstable repeat loci in our population.

## Methodology

### Sample enrolment

Cases - genetically uncharacterized SCA cases: 100 (Retrospective + Prospective Cases) (Autosomal dominant/ X-linked inheritance/ sporadic late age of onset patients), (negative for SCA1, SCA2, SCA3, SCA6, SCA7, SCA8, SCA12, SCA17 and FRDA. The mean age (SD), range of the patient was 59.07 (8.12), 42-84 years; the mean age of onset was 55.54 (7.11), 42-70 years.

Control - The control samples (N=100) were made available from the DNA repository of the Indian Genome Variation Consortium project (5). We divided the analysis into two stages (Fig 1), with the first focusing on finding the genome’s unstable CNG sites and the second trying to find out.

### A. In-silico identification of repeat loci in the human reference genome

The repeat sequences, CAG, CTG, CCG, CGG, and GCC which were reported for various ADCA sub-types were considered for this study (Fig 2). The program has been written in python to find all possible repeats with a minimum of 4 continuous repetitive units which are uploaded in the github. (https://github.com/bharathramh/STR_repeat/blob/main/str.py)

**Fig 2.**
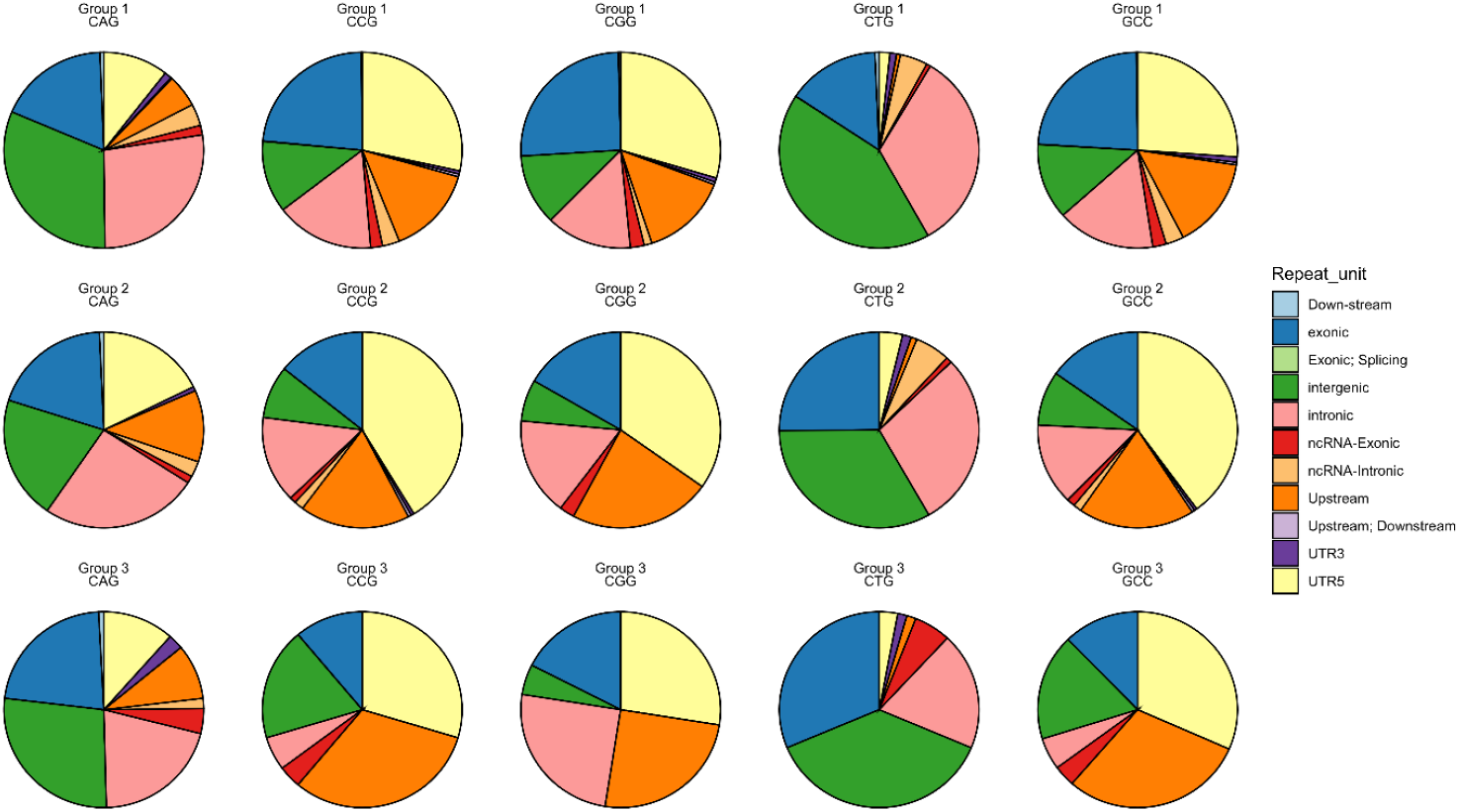
Classification of CNG repeat groups on the basis of functional regions in the genome

The python modules “Seq”, and “SeqIO” from the “Bio” package were used to read chromosome-wise fasta sequences and RegEx was generally used to find a sequence of strings in a specific pattern of repeats.

### Sub-Grouping of repeats on the bases of repeat number

The result of the program yielded 15069 repeats with more than 3 continuous repeat units. For better downstream analysis the repeats were duly categorized into three sub-types based on the length of the repeat (Table 1)

a. Group1 (4-6 repeats),
b. Group2 (7-9 repeats)
c. Group 3 (>9 repeats)

After classification of the repeats, to know the genomic feature the chromosome locations of each repeat were taken to get annotated. ANNOVAR (6) was used to annotate the gene information of each repeat. These annotated repeat groups were further classified based on their functional significance

**Table 1:**
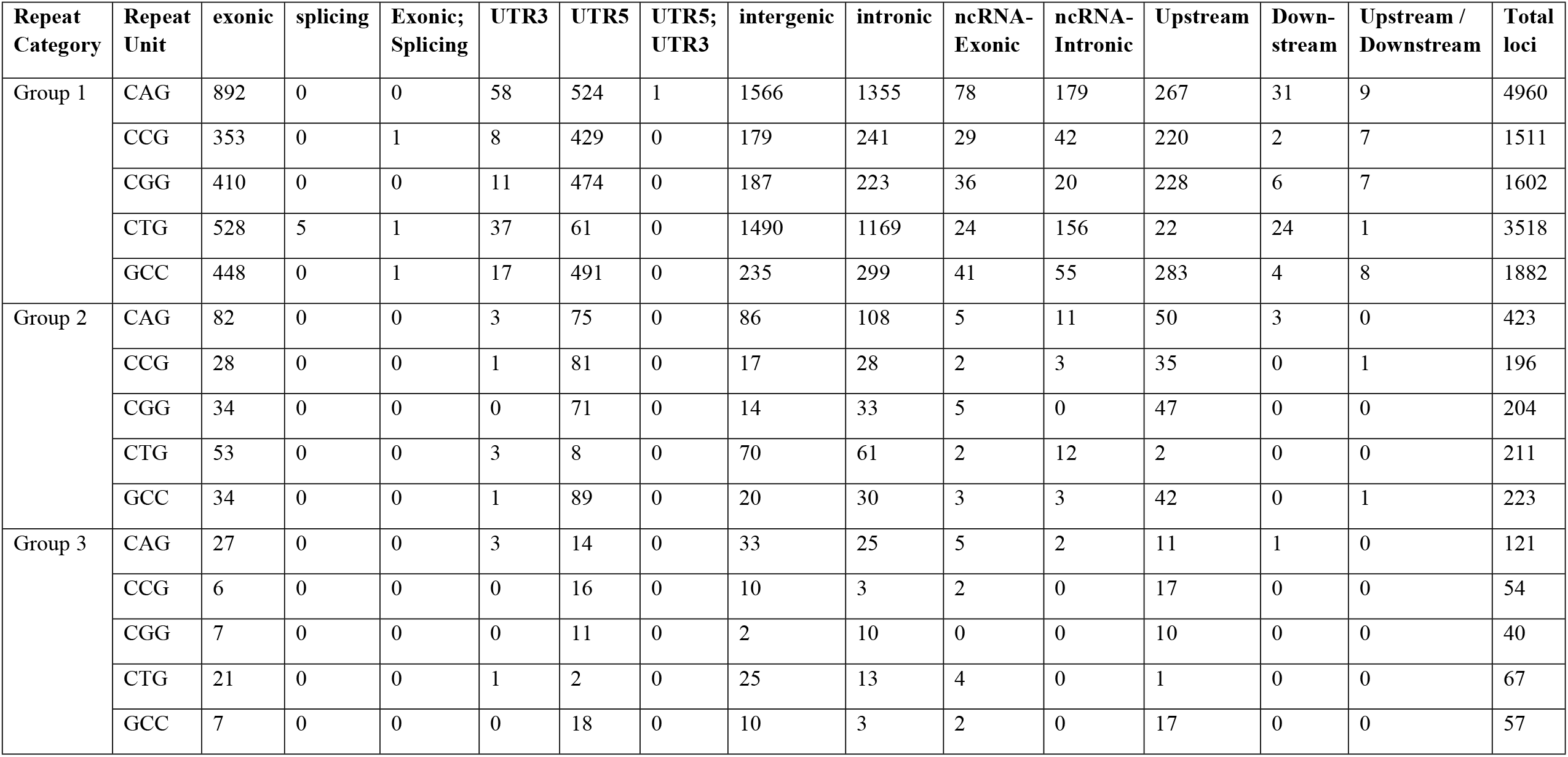
Classification of different CNG repeats on the basis of their size and functional domains

| Repeat Category | Repeat Unit | exonic | splicing | Exonic; Splicing | UTR3 | UTR5 | UTR5; UTR3 | intergenic | intronic | ncRNA-Exonic | ncRNA-Intronic | Upstream | Downstream | Upstream / Downstream | Total loci |
| --- | --- | --- | --- | --- | --- | --- | --- | --- | --- | --- | --- | --- | --- | --- | --- |
| Group 1 | CAG | 892 | 0 | 0 | 58 | 524 | 1 | 1566 | 1355 | 78 | 179 | 267 | 31 | 9 | 4960 |
|  | CCG | 353 | 0 | 1 | 8 | 429 | 0 | 179 | 241 | 29 | 42 | 220 | 2 | 7 | 1511 |
|  | CGG | 410 | 0 | 0 | 11 | 474 | 0 | 187 | 223 | 36 | 20 | 228 | 6 | 7 | 1602 |
|  | CTG | 528 | 5 | 1 | 37 | 61 | 0 | 1490 | 1169 | 24 | 156 | 22 | 24 | 1 | 3518 |
|  | GCC | 448 | 0 | 1 | 17 | 491 | 0 | 235 | 299 | 41 | 55 | 283 | 4 | 8 | 1882 |
| Group 2 | CAG | 82 | 0 | 0 | 3 | 75 | 0 | 86 | 108 | 5 | 11 | 50 | 3 | 0 | 423 |
|  | CCG | 28 | 0 | 0 | 1 | 81 | 0 | 17 | 28 | 2 | 3 | 35 | 0 | 1 | 196 |
|  | CGG | 34 | 0 | 0 | 0 | 71 | 0 | 14 | 33 | 5 | 0 | 47 | 0 | 0 | 204 |
|  | CTG | 53 | 0 | 0 | 3 | 8 | 0 | 70 | 61 | 2 | 12 | 2 | 0 | 0 | 211 |
|  | GCC | 34 | 0 | 0 | 1 | 89 | 0 | 20 | 30 | 3 | 3 | 42 | 0 | 1 | 223 |
| Group 3 | CAG | 27 | 0 | 0 | 3 | 14 | 0 | 33 | 25 | 5 | 2 | 11 | 1 | 0 | 121 |
|  | CCG | 6 | 0 | 0 | 0 | 16 | 0 | 10 | 3 | 2 | 0 | 17 | 0 | 0 | 54 |
|  | CGG | 7 | 0 | 0 | 0 | 11 | 0 | 2 | 10 | 0 | 0 | 10 | 0 | 0 | 40 |
|  | CTG | 21 | 0 | 0 | 1 | 2 | 0 | 25 | 13 | 4 | 0 | 1 | 0 | 0 | 67 |
|  | GCC | 7 | 0 | 0 | 0 | 18 | 0 | 10 | 3 | 2 | 0 | 17 | 0 | 0 | 57 |

### Selection of the repeat loci

The literature and our in-house data suggested that most of the ADCAs or late age of disease onset SCAs have been associated with polymorphic repeat expansion in coding and UTR regions. As the number of target loci is extremely high and unable to screen from conventional methods, we have focused on large CNG repeat loci (≥10 continuous repeats) associated with these regions for this study and finally selected 52 loci.

### B. Identification of unstable CNG repeat loci

To find out the polymorphic status in our population, we screened all the selected 52 CNG repeat loci in 100 control samples. To make our study cost-effective, we added an M13-specific nucleotide tag sequence on the 5’ end of all forward primers. Therefore, we used 3 primers for every PCR reaction (FP, RP, and fluorescent-labelled M13 tag primer).

For PCR amplification of selected 52 loci, we used different master mixes (Epicentre’s failsafe mixes or Promega master mix) along with 25 ng of DNA, 0.1 μl of forward primer, 0.4 μl of reverse primer, and 0.4 μl of M13 tag primer of 10 pM/μl working concentration in 10μl reaction volume. The PCR conditions were 95°C for 3 min followed by 35 cycles of denaturation at 95°C for 30 sec., annealing at 60°C for 30 sec, extension at 72°C for 30 sec, followed by a final extension at 72°C for 5 minutes. The samples were analysed using fragment analyser and visualized on gene mapper software (version 4, Applied Biosystems). After the repeat number calculation of all the loci and found 2 types of repeat status i) stable loci and ii) unstable loci (> ± 3 repeats variability). The highly polymorphic loci (Supplementary Table-1) were further selected for screening in patient samples.

## Results

From genome wide CNG repeat selection, we found a total of 15069 loci. The CNG repeats were found to abound in coding and UTR region. The CGG and CCG repeats were present mostly in the 5’UTR region of the gene (Fig-2).

We have screened all 52 loci with our control samples to know the repeat status in our population. Our data suggested, out of these, 33 loci were quite stable, and 19 loci were polymorphic in nature (Table-1). The unstable targets RAI1, UMAD1, GLS, HTR7P1, CNKSR2, MAML3, MED15, MLLT3, USF3, MEF2A, MIR205HG, NCOR2, RPL14, JPH3, MAB21L1, ANKUB1, ERF, GIPC1, and EP400 were further screened in the patient cohort to know any large repeat variability (Fig-3). The genes MAB21L1, ANKUB1, and GLS were highly polymorphic and had a wide range of repeat distribution in the population [Mode of repeats (Range): 13 (8 - 26); 15 (8 – 33); 12 (6 – 29) respectively]. The gene ANKUB1 and UMAD1 possess large normal repeats (>30 repeats) in both cases and control groups. No significant difference in large expansion range was observed in the case versus controls screening (Table 1). The heterozygosity indexes (HI) of UMAD1, MAB21L1, ANKUB1, GLS, and RPL14 show the repeat loci in these genes are highly polymorphic and greater than 0.7 both in cases and controls. On the other hand, MLLT3 and CNKSR2 were very less polymorphic and had more homozygous repeats (HI ≤ 0.1) in both groups. Most of the target loci fall between the range of 0.3 and 0.7 except ERF which also has a low HI of less than 0.25 in all samples.

**Figure 3:**
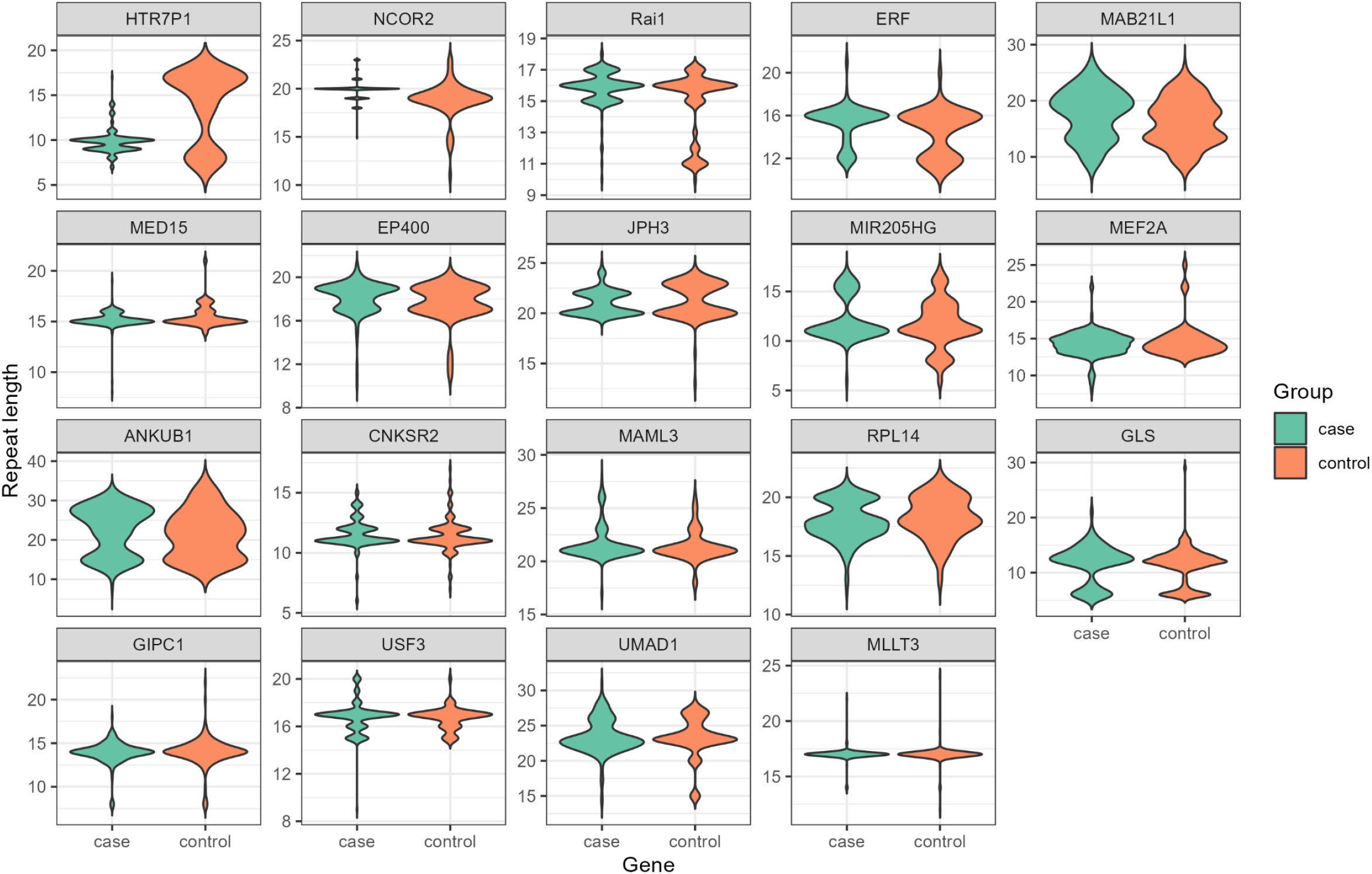
Repeat distribution of target repeats among control and patient samples

**Table 1:** Polymorphic status of selected loci in controls, patients, Heterozygosity index, 1000 genome database and GTEx brain expression

| Gene | Sample Group | Number of alleles | Repeat Range | Heterozygosity Index (HI) | Reference repeats number | Repeat status in 1000 G | Expression in Brain (GTEx) |
| --- | --- | --- | --- | --- | --- | --- | --- |
| GIPC1 | Controls | 11 | 8-22 | 0.27 | 10 | 6-22 | Present |
|  | Patients | 8 | 8-18 | 0.30 |  |  |  |
| UMAD1 | Controls | 9 | 15-28 | 0.62 | 20 | 4-24 | Present |
|  | Patients | 16 | 14-31 | 0.78 |  |  |  |
| MAB21L1 | Controls | 18 | 8-26 | 0.94 | 19 | 5-26 | Present |
|  | Patients | 18 | 8-26 | 0.91 |  |  |  |
| ANKUB1 | Controls | 14 | 14-33 | 0.85 | 20 | 6-26 | Present |
|  | Patients | 18 | 8-31 | 0.91 |  |  |  |
| GLS | Controls | 13 | 6-29 | 0.78 | 15 | 5-24 | Present |
|  | Patients | 12 | 6-21 | 0.89 |  |  |  |
| EP400 | Controls | 7 | 11-20 | 0.54 | 14 | 11-28 | Present |
|  | Patients | 11 | 10-21 | 0.50 |  |  |  |
| HTR7P1 | Controls | 6 | 8-17 | 0.61 | 10 | - | Present |
|  | Patients | 9 | 7-17 | 0.52 |  |  |  |
| RAI1 | Controls | 7 | 11-17 | 0.34 | 13 | 6-14 | Present |
|  | Patients | 9 | 10-18 | 0.46 |  |  |  |
| CNKSR2 | Controls | 11 | 7-17 | 0.21 | 12 | - | Present |
|  | Patients | 8 | 6-15 | 0.09 |  |  |  |
| MAML3 | Controls | 7 | 18-26 | 0.44 | 14 | 4-20 | Present |
|  | Patients | 9 | 17-28 | 0.34 |  |  |  |
| MED15 | Controls | 5 | 14-21 | 0.27 | 12 | 9-13 | Present |
|  | Patients | 7 | 8-19 | 0.33 |  |  |  |
| MLLT3 | Controls | 7 | 12-24 | 0.07 | 10 | - | Present |
|  | Patients | 5 | 14-22 | 0.05 |  |  |  |
| ERF | Controls | 9 | 11-20 | 0.23 | 10 | 6-11 | Present |
|  | Patients | 8 | 12-21 | 0.25 |  |  |  |
| USF3 | Controls | 5 | 15-20 | 0.52 | 11 | 6-16 | Present |
|  | Patients | 7 | 9-20 | 0.62 |  |  |  |
| RPL14 | Controls | 8 | 13-21 | 0.69 | 10 | - | Present |
|  | Patients | 10 | 12-21 | 0.79 |  |  |  |
| JPH3 | Controls | 6 | 13-24 | 0.61 | 15 | 5-19 | Present |
|  | Patients | 7 | 19-25 | 0.56 |  |  |  |
| MEF2A | Controls | 6 | 13-25 | 0.69 | 11 | 2-14 | Present |
|  | Patients | 10 | 8-22 | 0.61 |  |  |  |
| MIR205HG | Controls | 10 | 6-17 | 0.44 | 11 | 3-17 | Not Present |
|  | Patients | 7 | 6-17 | 0.46 |  |  |  |
| NCOR2 | Controls | 9 | 11-23 | 0.46 | 12 | 4-15 | Present |
|  | Patients | 7 | 15-23 | 0.51 |  |  |  |

### Data comparison of repeats among the different world populations

Since disease-associated tandem repeats tend to be more polymorphic in the general population, we plan to investigate the polymorphic nature of these loci in the control population. When compared with different 1000 genome control populations, the mode of repeats and variability in the GLS gene is higher in African and SAS populations (Table-2). MAB21L1 showed a repeat range on the higher side in the EAS population. Though some of the other loci had a maximum of >20 repeat expansions, they were uniform/less variable within the populations. MEF2A was highly variable from 2 to 16 repeats, but it is uniform throughout the population. GIPC1 repeat variability is less in the EUR population. For MED15 and ERF, repeat data is present of very few patient samples among different populations. We could not find any short tandem repeat data for HTR7P1, RPL14, CNKSR2, and MLLT3 repeat loci. our repeat data of GLS, ANKUB1, EP400, JPH3, and RAI1 loci showed bi-allelic distribution which is also seen in other major populations. Interestingly we observed a slightly large minimum and maximum repeat range in USF3, MEF2A, JPH3, RAI1, ERF, MED15, MAML3, and UMAD1 when compared to other world populations. Our both groups have comparatively fewer repeat numbers for EP400 loci. (Table 2)

**Table 2:** Repeat distribution of targeted repeat loci among different populations, our control and patient samples. *Repeat Mode (Minimum repeats – maximum repeats)

| S No. | GENE | AFR | AMR | EAS | EUR | SAS | Control | Case |
| --- | --- | --- | --- | --- | --- | --- | --- | --- |
| 1 | MIR205HG | 13 (4-18) | 9 (4-18) | 9 (3-14) | 9 (3-14) | 9 (4-16) | 11 (6-17) | 11 (6-17) |
| 2 | GLS | 14 (5-24) | 8 (5-23) | 8 (6-21) | 8 (6-22) | 14 (6-26) | 12 (6-29) | 12 (6-21) |
| 3 | USF3 | 13 (6-15) | 13 (7-17) | 13 (8-16) | 13 (6-16) | 13 (6-17) | 17 (15-20) | 17 (9-20) |
| 4 | ANKUB1 | 11 (8-25) | 11 (9-27) | 11 (6-26) | 11 (7-26) | 11 (6-26) | 15 (14-33) | 15 (8-31) |
| 5 | MAML3 | 18 (4-20) | 18 (8-19) | 18 (4-18) | 18 (6-19) | 18 (8-18) | 21 (18-26) | 21 (17-28) |
| 6 | UMAD1 | 18 (7-25) | 17 (6-25) | 17 (4-24) | 17 (6-24) | 17 (6-25) | 23 (15-28) | 23 (14-31) |
| 7 | NCOR2 | 17 (10-21) | 18 (13-22) | 18 (10-21) | 18 (4-22) | 18 (11-22) | 19 (11-23) | 20 (15-23) |
| 8 | EP400 | 28 (12-29) | 28 (12-29) | 28 (12-29) | 28 (11-29) | 28 (12-29) | 17 (11-20) | 19 (10-21) |
| 9 | MAB21L1 | 11 (5-27) | 13 (6-26) | 17 (4-26) | 13 (6-27) | 13 (5-26) | 13 (8-26) | 13 (8-26) |
| 10 | MEF2A | 11 (2-16) | 11 (2-13) | 9 (2-16) | 11 (3-13) | 9 (2-14) | 14 (13-25) | 15 (8-22) |
| 11 | JPH3 | 15 (6-26) | 15 (9-19) | 15 (10-19) | 15 (10-20) | 15 (7-25) | 20 (13-24) | 20 (19-25) |
| 12 | RAI1 | 12 (6-15) | 12 (7-15) | 12 (5-13) | 12 (6-14) | 12 (6-14) | 16 (10-17) | 16 (10-18) |
| 13 | GIPC1 | 13 (6-22) | 14 (9-17) | 14 (6-18) | 14 (12-20) | 14 (7-22) | 14 (8-29) | 14 (8-18) |
| 14 | ERF | 9 (6-10) | 9 (8-10) | 9 (9-9) | 9 (8-11) | 9 (6-11) | 16 (11-20) | 16 (12-21) |
| 15 | MED15 | 12 (12-12) | 12 (12-12) | 12 (12-12) | 12 (9-13) | 12 (11-12) | 15 (14-21) | 15 (8-19) |

The GTEx expression database (16) showed that genes CNKSR2, MAB21L1, USF3, RAI1, NCOR2, JPH3, MAML3, EP400, and GLS were significantly highly expressed in the brain, particularly the cerebellum. All the other genes were also showing significant expression in the brain except for MIR205HG (Table-1). Since SCA disorder is associated with the brain, we excluded MIR205HG from the shortlisted gene list; thus, we propose the pathogenicity of the remaining 18 genes which might show ataxia phenotype.

## Conclusion

The discovery of repeat instability is an underlying mutation mechanism for several neurodegenerative disorders in humans. It has always been a challenging task to understand the mechanism of repeat instability to disease manifestation. Several distinct hypotheses on the repeat expansion have been proposed over the years but still, its mechanism is not fully understood. Repeat instability in spinocerebellar ataxia contributes to the most prevalent genetic manifestation worldwide.

Our initial phase of the study through computational approach yielded 52 suspected CNG repeat loci from various genes for further investigation on the Indian control population. Using a cost-effective Fluorescent PCR based fragment analysis approach, resulted in 19 conclusive highly polymorphic repeat targets after screening in the control samples.

It has been proven that genetic markers for the same disorder are expressed among various populations in diverse ways. Hence few diseases and genetic markers are population specific.

So, in the second phase of the study, we screened these putative candidates in genetically uncharacterized clinically confirmed SCA samples. Although no large expansion of these target loci was identified in the study population. As a proof of concept, the repeat polymorphism in other populations of the 1000 genome data has been used. We tried to check all identified unstable markers in different major populations and our control and patient sample to know the population variability among these loci. We found repeat data of 15 loci out of 19 selected CNG loci from the 1000 genome STR database.

Alongside, the GTEx data showed that except MIR205HG, the remaining 18 loci had expression in various brain tissues which makes them more suitable for further investigations. In our study, we identified 18 highly unstable repeat loci but none of these showed large repeat expansion in our patient population.

Multiple studies published in recent years that were based on point and repeat expansion mutations for various neuro-related disorders from the proposed list of 18 genes, support our adopted strategy of the experiment of this study. It has been shown that the variation in the length of CAG repeats in the RAI gene is associated with age at onset modifier in spinocerebellar ataxia type 1 among various populations (7). In 2019 Rad et al. reported that point mutation in MAB21L1 causes a syndromic neurodevelopmental disorder with distinctive cerebellar, ocular, craniofacial, and genital features (COFG syndrome) (8). Another study suggested that point mutation in CNKSR2 is associated with seizures and mild intellectual disability (8). In 2020 a report was published hypothesizing that frameshift mutations of GLI3, ANKUB1, and TAS2R3 might alter the functions of proteins, and accelerate the progression of Polysyndactyly (PSD); an autosomal dominant genetic limb malformation (10). It has been proposed that the EP400 gene plays a significant role in oligodendrocyte survival and myelination in the vertebrate central nervous system (11). A study proposed that Gene MED15 polyglutamine repeat length changes the expression of diverse stress pathways (12). Among various populations, it has been reported that CAG repeat variation in MEF2A is a risk factor for coronary artery disease (CAD) (13). A study published in 2020 suggested that CGG repeat expansion mutation in 5’UTR of GIPC1 causes Oculopharyngodistal myopathy (OPDM) an adult-onset inherited neuromuscular disorder (13). A large GCA tandem expansion in 5’ UTR of the GLS gene causes overall development delay, progressive ataxia, and elevated levels of glutamine (14). Reported study of GLS, GIPC1, MED15, RAI1, and MEF2A has the same candidate loci that we identified in our study. All this reported evidence gives strength to our study although we did not find any large repeat expansion the approach is in the right direction in the discovery of a novel target.

### Limitations of the study

1. Repeat units: We considered only CNG repeats in coding and UTR regions with at least 10 continuous repeats due to screening hundreds of samples for a larger number of target loci. Considering other tri, tetra, penta, and hexa repeat units and lower repeat number loci increases the chances of getting causal mutations.
2. Lower sample size: We have taken 100 patient samples for the study. SCA is a rare type of disorder and its subtypes are very rare so considering a larger sample size will give more confidence to our hypothesis.
3. Unavailability of different population samples: It is well known that most of the SCA subtypes are geographic and population specific. In the study, we have considered only north Indian SCA patient samples where a multi-population study could enhance the possibility of finding causal mutation among studied genes.

This study highlights the importance of the population polymorphism approach to understand the genetic background and mechanism of tandem repeat instability in ataxia like neurological disorders. The role of other repetitive sequences in both coding and non-coding regions in context with neurological disorders can be explored with the help of computational and polymorphism approaches as taken in this work.

## Acknowledgement

We acknowledge the funding from CSIR-Young scientist project – OLP1127. And we acknowledge ICMR for the fellowship support for Varun Suroliya. We Thank UGC for the fellowship support for Manish Kumar. We thank CSIR-IGIB for the technical and scientific support. And also, Ataxia clinic, AIIMS for providing samples for this study. We thank sincerely to the patients and the families for their participation.

## Supporting information

**Supplementary table 1-** List of selected 52 loci along with repeat status in control samples (unstable loci marked in bold) and Primer details

